# Modular automated high-throughput isolation and phylogenetic identification of bacteria from complex microbiomes

**DOI:** 10.1101/2025.03.18.643884

**Authors:** Rubén Chaboy-Cansado, Silvia Talavera-Marcos, Ramón Gallego-Simón, Paula Cobeta, Gabriel Roscales, Alberto Rastrojo, Daniel Aguirre de Cárcer

## Abstract

Metagenomic analysis can generate hypotheses about microbiome interactions and function, yet mechanistic understanding is only possible through precise experimentation manipulating its microbiota composition. The high-throughput isolation of microbiome members thus represents a core resource in this field of research.

Here, we present and test a culturomics pipeline based on the use of the limiting dilution method with multi-well plates, an optical plate reader, and barcoded sequencing for phylogenetic identification, and offer modularity by proposing different protocols and possibilities along the pipeline. Most importantly, we provide all scripts required for process automation using an affordable pipetting robot, along with associated estimates of financial and labor costs.

## INTRODUCTION

Microbiomes are highly complex and extraordinarily diverse entities that control the Earth’s biogeochemical cycles and impact the biological fitness of their multicellular hosts. Our understanding of their genetic structure has greatly increased thanks to the development and democratization of high-throughput sequencing technologies. In this regards, metagenomic analysis can generate hypotheses about microbiome interactions and function, yet mechanistic understanding is only possible through precise experimentation manipulating its microbiota composition. Thus, the isolation of members of the resident microbiota represents a core resource for mechanistic studies aiming at dissecting microbiome interactions (1). Worldwide microbial collections and research labs now harbor thousands upon thousands of pure bacterial strains. Notably, some microbiomes, particularly human-associated microbiomes, have been extensively sampled through numerous small-scale studies as well as large-scale high-throughput projects (2).

Microbiome studies aiming to obtain mechanistic insights commonly start by assessing microbiota compositions using 16S rRNA phylogenetic marker gene sequencing or the more information-rich, yet labor-intensive and costly, shot-gun metagenomic sequencing. Subsequent bioinformatic analysis can generate hypotheses on the inner workings of the microbiome under study, which can be further tested using precise microbiota experimentation. To achieve this, the initial microbiome-derived sequences can be used to select strains available in microbial collections. However, it is unlikely that a set of microbes isolated from different, albeit similar, microbiomes would provide an accurate representation of the microbiota under study, as microbial adaptation is expected to contribute to genotypic and functional differentiation among microbes sampled from different habitats (1). Therefore, it is generally more appropriate to obtain a culture collection specific to the microbiome under study. Moreover, some hypotheses can be explored using genomic data from closely related genomes deposited in the databases or, preferably, using metagenomically-assembled genomes (MAGs) derived from the microbiome under study (e.g. investigating strain diversity (3) or modeling metabolic interactions using genome-scale models (4). However, genomes bearing exact or highly similar 16S rRNA gene sequences may exhibit substantial genomic heterogeneity (5), and reconstructing high-quality MAGs from complex communities still presents significant computational challenges (6, 7). Thus, direct isolation from the sample under study still represents the gold standard for describing genomic diversity (5).

The production of microbiome-representative culture collections through non-targeted microbial isolation is most commonly carried out using plating techniques on various solid media formulations and identifying selected colonies through Sanger sequencing of PCR-amplified 16S rRNA phylogenetic marker genes (e.g. (8)). Higher throughput can be attained by pre-filtering selected colonies using MALDI-TOF (9). However, these approaches have limitations. First, many strains may not be able to grow on solid media. Additionally, fast-growing bacteria may inhibit slow-growing ones, biasing the sampling effort towards fast growers. A common solution, picking colonies at different time points, is both labor-intensive and time-consuming. Sanger sequencing is expensive when considering increasing numbers of isolates and is inefficient at detecting contamination or co-isolation. The research group may not have access to (or experience with) a MALDI-TOF spectrometer or similar equipment. Finally, researchers often select colonies with different morphologies to enhance the diversity of their culture collection. However, this practice can be counterproductive when aiming for representativity; colonies with different morphologies are likely to pinpoint different underlying genomes, but most genomic diversity in a community is typically concentrated in a small number of shared morphology phenotypes.

Taking these limitations into consideration, Zhang *et al*. developed a high-throughput isolation protocol that can be carried out using common laboratory equipment, overcomes the challenges associated with colony picking, and leverages high-throughput sequencing of barcoded samples to streamline the process (1). Here, we present a pipeline based on their use of the limiting dilution method using multi-well plates and barcoded sequencing but improve upon it by employing 384-well plates to achieve higher throughput and incorporating an optical plate reader, available to most laboratories, to further reduce costs and increase throughput. Additionally, we introduce the possibility of using Oxford Nanopore sequencing which, for the purpose of high-throughput isolation, offers reduced costs, increased sensitivity and faster turnaround times. Moreover, we offer modularity by proposing different protocols and possibilities along the pipeline. Most importantly, we provide all scripts required for process automation using an affordable pipetting robot, along with associated estimates of financial and labor costs.

We conducted two consecutive tests for the proposed pipeline using tomato rhizosphere samples from our own research projects. First, we assayed the fully automated approach with complex microbiomes arising from tomato plants grown on different soils. Second, we processed three microbiome samples whose total diversity had been reduced for our own research needs using a multi-step *in planta* growth and dilution approach. In this case, we tested the fully automated approach against a hybrid pipeline that included an initial manual isolation step involving plating on solid medium and colony picking.

## MATERIALS AND METHODS

### Sample processing, DNA extraction, 16S rRNA gene sequencing and analysis of the original test samples

The root systems of three-week-old tomato plants were first shaken to remove loosely attached soil particles and then vortexed in 2 ml of a cold sterile 10 mM MgCl_2_ solution. For each rhizosphere fraction, 105 μl were frozen as 30% glycerol stocks at -20 ºC for less than a year before the subsequent isolation began. During the first test, total community DNA was extracted from 300 μl of each rhizosphere fraction using the *MagBind Environmental DNA 96 Kit* (Omega BioTek) according to the manufacturer’s instructions. For the second test, we processed 36 μl of each rhizosphere fraction following our automated version of Bramucci *et al*.’s alkaline lysis-based microvolume method (Bramucci et al. 2021). Using an automated two-step nested PCR approach, the 16S rRNA bacterial gene was initially amplified from the DNA samples using primers 341F and 805R, which target the V3-V4 hypervariable region, and including (3’-5’) a stretch of 0-7 Ns for frame shifting (Naik, Sharda, and Pandit 2020) and Illumina sequencing adapters (5’-TCGTCGGCAGCGTCAGATGTGTATAAGAGACAGN_0-7_CCTACGGGNBGCASCAG-3’ and 5’-GTCTCGTGGGCTCGGAGATGTGTATAAGAGACAGN_0-7_GACTACNVGGGTATCTAATCC-3’, respectively). Then, 1 μl of the resulting products was subjected to a second amplification using primers bearing (5’-3’) the required i5 and i7 Illumina adapters, 10 nt barcodes, and the 5’ end of Illumina sequencing primers (5’-AATGATACGGCGACCACCGAGATCTACACXXXXXXXXXXTCGTCGGCAGC GTC-3` and 5’-CAAGCAGAAGACGGCATACGAGATXXXXXXXXXXGTCTCGTGGGCTCGG-3’). Both PCR reactions consisted of a 44 µl reaction mixture containing 0.1 µM or 0.4 µM of each primer (first and second PCR steps, respectively), 0.4 mM of dNTPs, and 1 U of *Q5 HighFidelity* DNA Polymerase (New England Biolabs). The thermocycler conditions consisted of 95 °C for 30 s, followed by 20 or 10 cycles (first and second PCR steps, respectively) of 95 °C for 10 s, 55 °C for 30 s, and 72 °C for 30 s, with a final extension step of 2 min at 72 ºC. The amplicon libraries produced were checked using agarose gel electrophoresis and their concentration measured using *Picogreen* (Invitrogen). Equimolar amounts from each library were then pooled and run on an agarose gel. The appropriate-size band was gel excised and purified using the *QIAquick* Gel Extraction Kit (Qiagen). The final product was sequenced on an *Illumina MiSeq* NGS platform using a 600-cycle v3 reagent kit following the manufacturer’s instructions. Sequence processing was implemented in the *R* package *DADA2* (10) and included its standard pipeline for error modelling, paired-end sequence merging, chimera removal, taxonomic assignments using SILVA’s training set (v123) (11), and the elimination of residual eukaryotic sequences including mitochondria and chloroplast-affiliated sequences.

### Isolation and isolates processing

The approach to microbiome sample processing depends on the characteristics of the specific microbiome under study, and the detailed description of suitable protocols falls outside the scope of this article. In our case, we continued with the frozen glycerol stocks described above. The overall procedure starts by performing a preliminary cultivation experiment in which serial dilutions of the original sample are grown to assess the sample’s microbial load (Figure 1). The results are then used to calculate the desired dilutions for the isolation step. With the assumption that bacteria are distributed into different wells following a Poisson distribution, the predicted maximum number of wells containing clonal cultures is *ca*. 30% (12). To account for variability on the initial microbial load estimation and other experimental factors, we chose to produce four 384-well plates per run containing 1.5, 0.15 (two plates), and 0.015 bacteria per well, although proportions can be modified as desired. For both tests we used R2A medium, which is often employed to isolate soil bacteria. After an extensive period of two weeks of growth at 28ºC to allow for the isolation of slow growers (adapt temperature and time as needed), the OD600 of individual plates is recorded on a microplate reader (in our case a *BioTek Synergy HT reader)*. The results are then parsed to evaluate the suitability of the plate on the basis of the percentage of wells with growth (Zhang et al reported that plates containing 30-50% wells with growth generated a substantial proportion of pure cultures (1)) and to select the wells with above-threshold growth on each plate.

**Figure 1.**
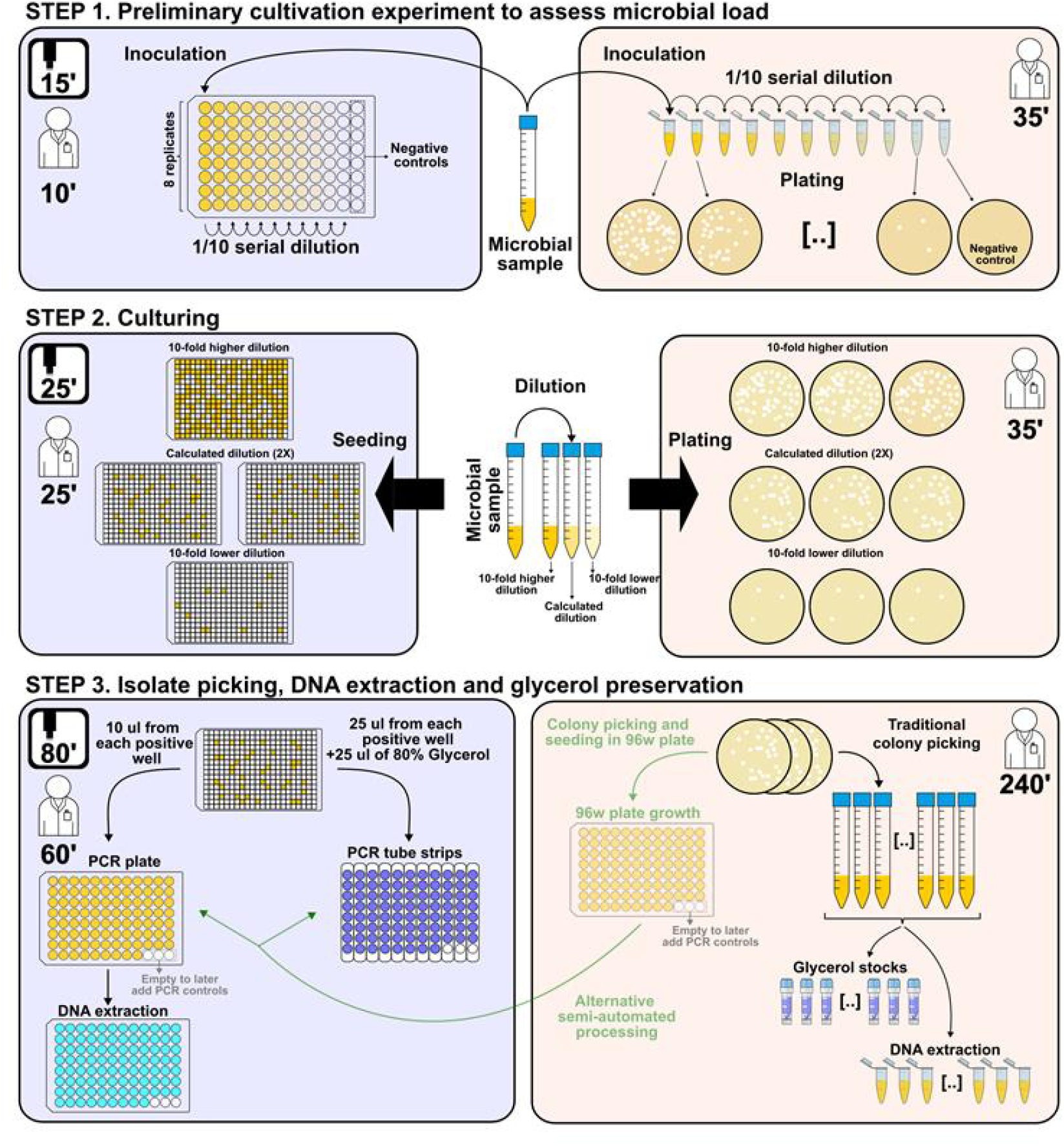
Diagram depicting the proposed automated pipeline and its manual counterpart (left and right, respectively). The diagram indicates the time needed to execute each step by researcher and robot. Additionally, step 3 shows the alternative automated processing of plate isolates (light green).

The automated procedure continues by processing the 384-well plate cultures; liquid samples from each selected well in a maximum of three 384-well plates are processed to produce a collection of up to 93 isolates per run. These samples are subsequently processed to produce temporal glycerol stocks and to be subjected to a standard alkaline lysis extraction in a single automated step. For the second test, we also isolated bacteria by plating appropriate dilutions on solid R2A agar plates. Single colonies were picked with a sterile pipette tip and used to inoculate 96 well culture plates with 200 μl of sterile R2A liquid medium. The plates were then incubated for 24 h at 28 ºC, and used to produce 30% glycerol stocks and extract DNA using our automated microvolume extraction method indicated above.

### Phylogenetic identification

Using the isolate lysates as template, we produced barcoded amplicon sequencing libraries following the same standard two-step nested PCR approach described above in 96 wells microtiter plates. However, in this case we chose to target the nearly full-length 16S rRNA gene by using the specific primer sequences 27f and 1492R during the first PCR. Also, we halved the reaction mixture volume and pooled per-plate amplicons based on agarose gel intensities to reduce costs. For the second PCR, wells in the same microtiter plate received a shared per-plate barcode and an individual per-well barcode. The advantage of this double barcoding strategy, featuring a shared barcode per plate and unique barcode per well, is that up to 9216 isolates can be analyzed in a single sequencing run using two sets of 96 barcoded primers. The per-plate amplicon pools were quantified using a *Qubit* spectrophotometer (Invitrogen), mixed in equimolar proportions and the appropriate-size band was gel excised and purified as above.

The resulting amplicon pool was transformed into a sequencing library using the Ligation Sequencing DNA V14 kit (Oxford Nanopore) and sequenced using a *MinION* sequencer (Oxford Nanopore) with a 10.4.1 flowcell. Nevertheless, the two-step PCR approach was performed using primers prepared for *Illumina* sequencing, so exactly the same procedure can be undertaken with *Illumina* sequencing albeit with primers targeting a smaller region (e.g. primer sequences 341F and 805R and a 2 x 300 bp sequencing).

For the processing of *Nanopore*-derived sequences, we provide scripts to demultiplex the raw data and generate per well consensus sequences along with a quality indicator to evaluate contamination/co-isolation. For the processing of *Illumina* paired-end sequences we recommend the excellent purpose-built bioinformatic pipeline (github.com/YongxinLiu/Culturome) developed by Zhang et al (1).

As a final additional step, chosen isolates can be recovered from their glycerol stocks and further purified re-streaking the isolates a number of times on agar plates, producing a new glycerol stock and re-sequencing. For this purpose, there are two straightforward possibilities: i) long 16S rRNA gene PCR, followed by magnetic beads-based purification and Sanger sequencing, or ii) barcoded high-throughput sequencing using the same procedure described above.

## RESULTS AND DICUSSION

### Isolation results

To test the ease and trustworthiness of the approach and automated protocols, we followed two different scenarios, with each different step carried out by two different researchers. In the first scenario, we carried out a single isolation run for each of eight tomato rhizosphere samples arising from different soils. In this case, we were able to obtain 95 pure isolates with different 16S rRNA gene sequences, spanning seven of the ten most abundant bacterial families present in the original samples (no isolates were obtained for the abundant Chitinophagaceae, Bacillaceae and Streptomycetaceae families). In the second scenario, we processed three microbiome samples bearing reduced diversities (16, 15, and 10 bacterial ASVs with a relative abundance > 0.8%). After the isolation and sequencing of a single batch of 478 clones (218 and 260 processed using the fully-automated or hybrid pipeline, respectively), we were able to recover all but one of the ASVs as isolates (> 97%).

### Time and costs

Although protocol run times and associated costs will vary to some degree from lab to lab, here we provide estimates grounded on protocol executions from two researchers and local consumable pricings as references (see Supplementary information). In addition to basic molecular biology and microbiology equipment, the full automation of the proposed pipeline requires the use of a microtiter plate reader, as well as an *Opentrons* OT-2 pipetting robot with multiple pipettors, HEPA filter, and thermocycler modules. At the time of writing, these robot parts added to 29,000$, which would go up to 32,750$ if the magnetic module is added for the proposed automated *Sanger* sequencing script, or drop to 19,250$ if an off-robot thermal cycler is used. These sums represent a significant investment; hence, we suggest those groups wishing to use the proposed pipeline in very few occasions and with rather shallow isolation needs to employ the proposed manual alternatives. On the other hand, for groups, such as ours, where high-throughput isolation from microbiomes is a core procedure, or research platforms and companies wishing to incorporate high-throughput isolation capabilities to their list of services, such initial investment would be justified. In addition to the previously mentioned follow-up mechanistic studies of microbiome patterns (e.g. (13)), a non-exhaustive list of fields likely requiring frequent high-throughput isolation procedures would surely include experimental evolution of microbial communities (14) and the search for plant growth promoting bacteria (8), among others. Significantly, the OT-2 is a flexible and accessible liquid handler that can automate a very large number of molecular biology and microbiology processes, such as the recent development of automated low-cost whole-genome sequencing of microbiota (5), more so with the proposed suit of modules, and thus its acquisition should be contemplated as a general lab investment not merely as a need to conduct high-throughput isolation from microbiomes.

### Manual approach

All steps of the pipeline can also be performed manually with the optional use of multichannel pipettes. However, this approach requires more time and carries a significantly higher risk of human error. If working manually with 384-well plates proves too cumbersome, the procedure can be adapted to 96-well plates. In the absence of a plate reader, microbial growth can be assessed visually by inspecting the wells. Later in the pipeline, the only equipment required includes a thermal cycler and agarose electrophoresis equipment, alongside standard molecular biology techniques. Finally, *Sanger, Oxford Nanopore*, and *Illumina* sequencing are widely available through numerous companies worldwide, should in-house resources be unavailable. Additionally, if the research team has access to a different robotic workstation, all or part of the pipeline may be executed using it, depending on its capabilities.

### Sequencing platform

The choice of sequencing platform for phylogenetic identification involves a trade-off between several factors; *Sanger* sequencing could be employed in the first screening round following the provided protocols, although its cost increases linearly with the number of isolates, since the methodology is refractory to barcoding strategies and hence each isolate would have to be sequenced individually. Nonetheless, it may be a suitable option for subsequent re-isolation screens if a low number of selected isolates makes the fixed costs of high-throughput sequencing too burdensome. High-throughput sequencing technologies, on the other hand, can incorporate barcoding strategies, significantly reducing sequencing costs if the number of isolates to be analyzed is large enough to offset their fixed costs. With sufficient sequencing depth per barcode, *Oxford Nanopore* sequencing achieves the same resolution as *Sanger* sequencing, as both technologies are capable of sequencing nearly full-length 16S rRNA gene amplicons. Compared to *Illumina, Oxford Nanopore* provides lower fixed costs and faster turnaround times if in-house *MiniION* sequencing is available and reusable flowcells or inexpensive low-throughput *Flongle* flowcells are used for shallow sequencing. Finally, *Illumina* sequencing can be employed for ultra-high throughput albeit reduced resolution due to the shorter read length.

### Limitations and modifications

The protocols provided here can be executed with basic training in microbiology and molecular biology techniques. However, familiarity with the Opentrons API is required for automation, and basic knowledge of Unix is necessary to run the sequence processing scripts and manage dependencies downloads. The current approach, in its present form, is suitable only for isolating bacteria able to grow in a chosen liquid medium. Nonetheless, the molecular biology component of the pipeline, along with the provided off-pipeline scripts, can be applied for the high-throughput phylogenetic identification of bacteria that are manually isolated on solid medium. This molecular biology component can also facilitate the high-throughput characterization of anaerobes, provided their culturing requirements are met manually. Alternatively, an OT-2 robot could be adapted for use within a sufficiently large anaerobic chamber, though interactions between the researcher, the chamber, and the robot -such as changing modules and pipettes or loading labware and media-may prove cumbersome. Matrix assisted laser desorption ionization-time of flight (MALDI-TOF) mass spectrometry has been used for the high-throughput characterization of bacterial isolates (15). Its integration with the proposed pipeline would be feasible, provided a suitable high-throughput protocol is developed to enable the transfer of dense liquid cultures to MALDI target spots and their proper preparation ana analysis. In this regard, the OT-2’s ability to incorporate custom labware would facilitate the spotting of liquid samples onto target spots. However, assessing whether the necessary physical and chemical processing of the liquid samples (e.g. without centrifugation) could be effectively achieved with the proposed platform falls beyond the scope of this work.

The targeted isolation of specific microbiome members has been achieved using fluorescence-activated cell sorting (FACS) in combination with various labelling approaches. These include targeting DNA (Live-FISH), incorporated metabolites (Metabolite fluorescent labelling or isotope labeling and Raman detection), or using target-specific antibodies (for a review see (16)). All these methods can be viewed as sample enrichment strategies that can be used jointly with the proposed pipeline, with the understanding that only pre-selected bacteria able to grow on the chosen medium would be isolated and characterized. Similarly, FACS-free antibody-based immunomagnetic precipitation (17) could be used to process a sample prior to using the proposed pipeline, allowing fOr the depletion or enrichment of the sample for specific bacteria before isolation.

### Conclusion

In this work, we have improved Zhang *et al*.’s original high-throughput isolation protocol by employing 384-well plates and incorporating an optical plate reader to achieve higher throughput. Also, we offer modularity with different protocol possibilities along the pipeline. Most importantly, we offer pipeline automation using an affordable pipetting robot. Recently, Huang et al (18) published an image-guided machine-learning directed high-throughput approach that picks colonies from agar plates based on their characteristics. By housing their system within an anaerobic chamber and employing a picking strategy targeting the “more diverse” colonies, they successfully isolated a significant portion of the diversity in human fecal samples. In addition to avoiding the issues related to solid-medium isolation, our automation solution is approximately nine times less expensive and, importantly, can be directly applied to automate other common microbiology and molecular biology tasks. Therefore, considering its modular design and cost-effectiveness, our proposed pipeline provides high-throughput isolation capabilities accessible to the broader Microbiome research community.

## Data availability

All scripts, companion protocol guides and sequences are available at github.com/microenvgen/Culturomics.

## Funding

This work was funded by the Spanish Ministry of Science and Innovation grant TED2021-130616B-I00 awarded to DAC.

## Conflict of interest

The authors declare no conflicts of interest.

## Notes

https://github.com/microenvgen/Culturomics

### Competing Interest Statement

The authors have declared no competing interest.

