## Supplementary information for "Modular automated high-throughput isolation and phylogenetic identification of bacteria from complex microbiomes"

The present document provides information on the materials, costs, time and troubleshooting associated with the proposed culturomics pipeline in both its automated and manual versions. For the manual version, the time estimates assume that the researcher uses a multichannel pipette.

**STEP 1. Preliminary cultivation experiment to assess microbial load**

**Automated pipeline:**

**Materials:** P200 tips, 96-well culture plate, suitable growth medium, 1.5ml polypropylene tubes.

**Total cost:** 2.22 euro.

**Troubleshooting:** All or most wells show microbial growth: the liquid medium was contaminated or HEPA filter malfunction. No microbial growth: unsuitable culture medium or no live bacteria in sample.

**Time:** Researcher; 10 min. Robot; 15min.

**Manual pipeline:**

**Materials:** P200 tips, petri dishes, suitable growth medium, 1.5ml polypropylene tubes.

**Total cost:** 1.5 euro (aprox).

**Troubleshooting:** Lack of growth on petri dishes: inadequate medium or no live bacteria on sample or wrong serial dilution. Too much growth on plates to count: adjust serial dilution. Incongruent results between plate counts and serial dilutions: wrong serial dilution.

**Time:** 35 min.

**STEP 2. Culturing**

**Automated pipeline:**

**Materials:** P200 tips, 384-well culture plates, suitable growth medium, polypropylene deposit.

**Total cost:** 38.6 euro.

**Troubleshooting:** All or most wells show microbial growth: the liquid medium was contaminated or HEPA filter malfunction or wrong sample dilution preparations (if negative controls are blank). No microbial growth: unsuitable culture medium or no live bacteria in sample or wrong sample dilution preparations.

**Time:** Researcher; 25 min, Robot 25; min.

**Manual pipeline:**

**Materials:** P200 tips, petri dishes, suitable growth medium, 1.5ml polypropylene tubes.

**Total cost:** 1.5 euro (aprox).

**Troubleshooting:** Lack of growth on petri dishes: inadequate medium or wrong serial dilution. Too much growth on plates: wrong serial dilution (adjust).

**Time:** 25 min.

**STEP 3. Isolate picking, DNA extraction and glycerol preservation**

**Automated pipeline:**

**Materials:** P200 tips, 96-well PCR plate, 0.2ml polypropylene tube strips, polypropylene deposit, glycerol and chemical reagents.

**Total cost:** 11.4.

**Troubleshooting:** NA

**Time:** Researcher; 60 min., Robot; 80 min.

**Remarks:** Visually inspect a few wells with above-threshold OD600 values to make sure that the chosen threshold indeed represents microbial growth.

**Manual pipeline:**

**Materials:** P200 tips, culture medium, 96-well culture plate, 0.2ml polypropylene tube strips, 96-well PCR plate, polypropylene deposit, glycerol and chemical reagents.

**Total cost:** 12 euro (approx).

**Troubleshooting:** NA

**Time:** 240 min during a two to three-days period (depending on the growth rate of the isolates).

**STEP 4. Amplicon library production (PCRs)**

**Automated pipeline:**

**Materials:** P200 tips, P20 tips, 96-well PCR plate, 15ml polypropylene tube, plate adhesive seal, PCR reagents.

**Total cost:** 186.2 euro.

**Troubleshooting:** In addition to standard PCR troubleshooting: if too many reactions fail, then the DNA extraction failed (check reagent pHs and reagent order in deposit during previous step) or the OD600 threshold set was too low or most isolates are resistant to DNA extraction approach.

**Time:** Researcher; 40 min. Robot; 12 min (not including the PCR itself).

**Manual pipeline:**

**Materials:** P200 tips, P20 tips, 96-well PCR plate, 15ml polypropylene tube, plate adhesive seal, PCR reagents.

**Total cost:** 186.2 euro.

**Troubleshooting:** In addition to standard PCR troubleshooting: if too many reactions fail, then the DNA extraction failed (check reagent pHs) or most isolates are resistant to DNA extraction approach.

**Remarks:** Take extra care not to mistake the wells.

**Time:** Researcher; 31 min.

**STEP 5. Pooling of amplicon libraries**

**Automated pipeline:**

**Materials:** P20 tips,1.5 ml polypropylene tube.

**Total cost:** 7.9 euro.

**Troubleshooting: T**he final volume exceeds the capacity of the polypropylene tube: wrong normalization file or error during the parsing of the file.

**Time.** Researcher; 25 min. Robot; 51 min.

**Manual pipeline:**

**Materials:** P20 tips,1.5 ml polypropylene tube.

**Total cost:** 7.9 euro.

**Troubleshooting:**

**Remarks:** Take extra care not to mistake the wells.

**Time.** Researcher; 55 min.
